# Deletion of CD44 promotes adipogenesis by regulating PPARɣ and cell cycle-related pathways

**DOI:** 10.1101/2023.10.13.562173

**Authors:** Xiong Weng, Hao Jiang, Houjiang Zhou, De Lin, Jing Wang, Li Kang

## Abstract

CD44, a cell surface adhesion receptor and stem cell biomarker, is recently implicated in chronic metabolic diseases. Ablation of CD44 ameliorates adipose tissue inflammation and insulin resistance in obesity. Here, we investigated cell type specific CD44 expression in human and mouse adipose tissue and further studied how CD44 in preadipocytes regulates adipocyte function. Using Crispr Cas9-mdediated gene deletion and lentivirus-mediated gene re-expression, we discovered that deletion of CD44 promotes adipocyte differentiation and adipogenesis, whereas re-expression of CD44 abolishes this effect and decreases insulin responsiveness and adiponectin secretion in 3T3-L1 cells. Mechanistically, CD44 does so via suppressing *Pparg* expression. Using quantitative proteomics analysis, we further discovered that cell cycle-regulated pathways were mostly decreased by deletion of CD44. Indeed, re-expression of CD44 moderately restored expression of proteins involved in all phases of the cell cycle. Our data suggest that CD44 plays a crucial role in regulating adipogenesis and adipocyte function possibly through regulating PPAR☐ and cell cycle-related pathways. This study provides evidence for the first time that CD44 expressed in preadipocytes plays key roles in regulating adipocyte function outside immune cells where CD44 is primarily expressed. Therefore, targeting CD44 in (pre)adipocytes may provide therapeutic potential to treat obesity-associated metabolic complications.

## 1. Introduction

Obesity is a metabolic complication of excess fat accumulation in organs, including adipose tissue, liver, skeletal muscle, and pancreas. The increasing obesity epidemic contributes to the prevalence of obesity-associated metabolic diseases, such as type 2 diabetes (T2D), non-alcoholic fatty liver disease and cardiometabolic conditions (Klein et al., 2022, Stefan, 2020). Over 670 million adults are obese worldwide, which causes huge socioeconomic burden to our society (Collaboration, 2017). Despite great efforts being made into elucidating the pathogenesis of obesity, mechanisms of the development of obesity are not fully understood. Thus, it’s important and urgent to identify novel therapeutic targets to advance the prevention and treatment of obesity and obesity-associated metabolic complications.

CD44 is a membrane protein and has different variants rising from alternative splicing in the *Cd44* gene (Ponta et al., 2003). The standard form of CD44 (CD44s) without alternative splicing variants is the smallest CD44 isoform (Zöller, 2011). CD44s is ubiquitously expressed in a wide variety of tissues including the central nervous system, liver, lung, adipose tissue and muscle (Weng et al., 2022). CD44 was intensively studied in cancer initiation and tumorigenesis, where increased expression of CD44 is often found as a biomarker of advanced tumour progression and poor prognosis in many cancers (Zöller, 2011, Hassn Mesrati et al., 2021). However, recent studies suggest a potential role of CD44 in chronic metabolic diseases, such as obesity and T2D (Weng et al., 2022). Increased CD44 expression and deposition of its ligands - hyaluronan (HA) and osteopontin (OPN), were found in the adipose tissue of obese mice and humans (Kodama et al., 2012). The plasma CD44 level was positively correlated with insulin resistance and poor glycaemic control in human (Liu et al., 2015). Moreover, *Cd44* gene was implicated in the pathogenesis of T2D by a gene expression-based genome-wide association study (Kodama et al., 2012). Ablation of CD44 by genetic deletion or anti-CD44 monoclonal antibody administration reduced adipose tissue immune cell infiltration and insulin resistance in diet-induced obese mice (Kodama et al., 2015). These studies highlight that CD44 has critical roles in regulating adipose tissue function during obesity. Adipocytes are essential in maintaining adipose tissue function, and adipogenesis and insulin response are key characteristics of adipocyte function. However, how CD44 regulates adipogenesis and insulin signalling in adipocytes, therefore contributing to adipose tissue function is unclear.

In this study, we utilised CRISPR Cas9-mediated gene deletion and lentivirus-mediated gene re-expression in 3T3-L1 cells to determine the role of CD44 in adipogenesis and insulin response in adipocytes. Mechanistically, quantitative proteomics analysis was used to explore potential pathways regulated by CD44 in preadipocytes. We found that deletion of CD44 promoted adipogenesis and insulin response in 3T3-L1 cells, which were mediated by upregulation of *Pparg* and changes in cell cycle-related pathways.

## 2. Materials and methods

### 2.1 3T3-L1 cell culture, differentiation, and insulin treatment

3T3-L1 cells were maintained in basal medium (4.5g/L high glucose DMEM medium, 10% Bovine Calf Serum, 1% Pen/Strep antibiotics, 2mM L-Glutamine and 5% 20mM HEPES). Cells were differentiated in medium containing 167nM insulin, 0.5mM 3-isobutyl-1-methylxanthine (IBMX), 1μM Dexamethasone (Dex) and 2μM Rosiglitazone (RAZ) for 4 days, followed by basal medium containing 167nM insulin for another 3 days. Cells were then maintained in basal medium for another 7 days. Unless indicated otherwise, all experiments were performed on Day 14 after differentiation initiation. For insulin treatment, cells were incubated with 100nM insulin for 30mins.

### 2.2 Oil Red O staining

Adipogenesis was assessed by Oil Red O staining. Cells were fixed in 4% formaldehyde for 2hrs before being rinsed with 60% isopropanol and subsequently stained with Oil Red O solution (0.3% w/v) for 20mins at room temperature. Staining was quantified by measuring the absorbance at 540nM.

### 2.3 qRT-PCR

Total RNA was extracted by Trizol reagent and synthesized into cDNA by SuperScript™ II Reverse Transcriptase kit (#18064014, ThermoFisher Scientific). qRT-PCR was run by either Taqman or SYBR^TM^ Green assays. The TaqMan probes for *Cd44* (Mm01277161_m1), *Pparg* (mm00440940_m1), *Cebpa* (Mm00514283_s1) and *18s* (Hs99999901_s1) were obtained from ThermoFisher. The primers of *Pref-1*: 5’-GGATTCGTCGACAAGACCTG-3’, 5’-GCTTGCACAGACACTCGAAG-3’; *Npr3*: 5’-GCAAATCATCAGGTGGCCTA-3’, 5’-CCATTAGCAAGCCAGCACCTA-3’; *Scara5*: 5’-TGTGGAAGGTTCAGGATGCG-3’, 5’-GGCTT CGATT GCTTTCCACC-3’; and *18s*: 5’-GCAATTATTCCCCATGAACG-3’, 5’-GGCCTCACTAAACCATCCAA-3’ were obtained from Sigma. Data were normalized to 18s and analysed using the 2^-ΔΔCt^ method.

### 2.4 Western blot

1x10^6^ cells or ∼50mg tissues were homogenised in protein lysis buffer (25mM Tris-HCI pH 7.4, 50mM NaF, 0.1mM NaCI, 1mM EDTA, 5mM EGTA, 9.2% sucrose, 1% Triton X100, 10mM NaPp, 0.1% Mercaptoethanol, 1mM Na_3_VO_4_, 1mM Benzamidine, 0.1M PMSF and 10% glycerol). Protein concentration was quantified, and proteins were separated by the SDS-PAGE gel before being imaged by Western blotting. The primary antibody for CD44 (#A303-872A-M, BETHYL Laboratories), AKT (#9272, Cell signalling), pAKT(S473) (#9212, Cell signalling), and β-tubulin (#ab6046, Abcam) were used at 1:1000 dilution. The secondary antibody: anti-rabbit (#P/N:926-32213, LI-COR) and anti-sheep (#NL010, R&D) were used at 1:100000 dilution.

### 2.5 Crispr-Cas9 gene editing

Guide RNAs (gRNAs) targeting mouse CD44 (NM_001039151.1) were designed by Broad Institute Portal (http://www.broadinstitute.org/rnai/public/analysis-tools/sgrna-design). The top 3 gRNAs that target early exons with maximal on-target effects were selected and cloned into PX459 plasmids. Undifferentiated 3T3-L1 cells were transfected with lipofectamine LTX (#A12621; ThermoFisher Scientific) for 48hrs, before being selected with 4μg/ml puromycin. The Crispr-Cas9 editing efficiency was assessed by CD44 protein expression by Western. Cell populations with the least CD44 expression were sorted into single cells by fluorescence-activated cell sorting (FACS). The single cell derived stable CD44 knockout (KO) cell lines were further validated by CD44 protein expression and characterised by bi-allelic sequencing. For bi-allelic sequencing, genomic DNAs of CD44KO cell lines were isolated and DNA sequences around the gRNA targeted site were amplified by PCR using primers 5’-GTGGTAATTCCGAGGATTCA-3’ and 5’-GGCTGTTCATGGCTGTTC-3’. The PCR products were cloned into a pSC-AMP/Kan vector, using the StrataClone PCR Cloning Kit (#240205, Agilent). The bi-allelic sequencing was performed with the M13 reverse primer: 5’-GAG CGG ATA ACA ATT TCA CAC AGG-3’.

### 2.6 Lentivirus-mediated CD44 re-expression

The CD44s DNA coding sequence (NP_001034240.1) was synthesized by Sigma and constructed into a lentivirus expressing vector *pLenti-CD44-C-mGFP-P2A-Puro* with a GFP tag. Both CD44 expressing vector and the control vector were packaged in HK293T cells, using the package plasmids PSPAX2 and PMD2.G with lipofectamine p3000 (#L3000075, ThermoFisher Scientific). CD44-expressing and control lentiviruses were then used to infect CD44KO 3T3-L1 cells for 48hrs. CD44KO 3T3-L1 cells were selected by 4μg/ml puromycin for 72hrs and the infection efficiency was validated by Western blot or qRT-PCR.

### 2.7 Sample preparation for proteomics analysis

Undifferentiated CrisprWT EV, CD44KO2 EV and CD44KO2 RE cells were prepared for proteomics analysis, following the single-pot solid-phase-enhanced sample preparation protocol (Hughes et al., 2019). Briefly, 1x10^6^ cells were lysed with 4% SDS in 100mM TEAB buffer (#90114, ThermoFisher Scientific) and denatured at 95[for 10mins. The cell lysates were then sonicated for 30 cycles (30s on, 30s off) before being treated with 10mM DTT for 1h for the reduction of reversibly oxidized cysteines. The proteins were then treated with 20mM iodoacetamide for 45mins at room temperature to alkylate free thiols. The protein concentration was quantified and 100ug proteins of each sample were incubated in acetonitrile with Sera-Mag SpeedBead Carboxylate-Modified Magnetic Particles (Hydrophilic) (#GE44152105050250, Merck) and Sera-Mag SpeedBead Carboxylate-Modified Magnetic Particles (Hydrophobic) (#GE24152105050350, Merck) for 10mins at room temperature. The beads were then washed and redissolved in 50mM ammonium bicarbonate and digested overnight with trypsin at 37°C. After being acidified with 10% formic acid, the peptides were washed and eluted in 100µL 2% DMSO. The samples were centrifuged at 10,000g and the peptide containing supernatants were dried by speedvac. The peptides were redissolved in 100mM TEAB buffer and equal amounts of peptides from each sample were labelled with TMT 10-plex Mass Tag labelling kit (#90110, ThermoFisher Scientific). After labelling, all samples were pooled and desalted before the mass spectrometry analysis.

### 2.8 LC-MS and data analysis

MS analysis of TMT-labelled peptides was performed on a Q-exactive-HF mass spectrometer coupled with a Dionex Ultimate 3000RS (ThermoFisher Scientific). The detailed LC-MS protocol can be found in the supplemental materials. The LC-MS raw data were searched against the IPI mouse database version 3.83 using MaxQuant (1.6.6.0). The corrected reporter ion intensity of each protein intensity ratio was used for subsequent analysis; the protein intensity results of each group were processed by Perseus (2.0.6.0). All data were transformed into log, regrouped according to genotypes and normalized to median. Unpaired student t-test was performed to detect significantly changed proteins. Differentially expressed proteins (DEPs) were defined as proteins with a fold change > 1.5 and a *p* value < 0.05. DEPs were visualized as volcano plots using Prism and the pathway enrichment was analysed by metascape.

### 2.9 Single cell sequencing data analysis

CD44 clustering expression in the subcutaneous and visceral white adipose tissue (WAT) of human and mice was analysed using published single cell RNA sequencing data (SCP1376) at Single Cell Portal (https://singlecell.broadinstitute.org/). Briefly, 363,870 cells with 166,149 of human cells and 19,7721 of mice were analysed according to their original clustering (Emont et al., 2022). The human single-cell data were integrated from paired visceral and subcutaneous WAT of plastic surgery biopsies of ten subjects and subcutaneous WAT from three adult males. The mouse WAT single-cell data contained gene expression profile of 197,721 cells, which integrated mouse subcutaneous and epididymal WAT of both sexes (10 males and 4 females) (Emont et al., 2022). For CD44 clustering expression in the epididymal WAT of obesity, single-cell RNA-seq data were derived from another public database (SCP1179) of male C57BL/6 mice fed either low fat diet (LFD) (Research Diets, 10% fat, D12450B) or high fat diet (HFD) (Research Diets, 60% fat, D12492) (Sárvári et al., 2021). 10X Genomics Chromium platform was used to sequence single-nuclear RNA-Sequence. A total of 19,723 cells were detected and clustered into seven different adipocyte subpopulations, including adipocytes (n=4604), endothelial cells (n=110), epididymal cells (n=499), adipose stem and precursor cells (ASPCs) (n=5171), immune cells (n=8354), mesothelial cells (n=680), and Spermatozoa (n=305). We extracted *Cd44* gene expression from each cell and performed differential expression analysis in different subpopulations using Seurat (PMID: 31178118).

### 2.10 Statistical analysis

All data were confirmed to be normally distributed and analysed by either unpaired student t-test or two-way ANOVA for statistical significance as indicated. Data were presented as mean±SEM and the significant level was *p<0.05, **p<0.01, ***p<0.005, and ****p<0.001. All data figures were generated by Prism (Graphpad).

## 3. Results

### 3.1 CD44 expression was regulated by obesity and during adipocyte differentiation

CD44 protein was increased by 5 folds in the epididymal WAT of HFD-fed male mice compared to those of chow-fed males (Figure 1A and B). The expression pattern of CD44 in mouse and human WAT was next analysed using published single cell sequencing data (Emont et al., 2022). It was shown that CD44 was ubiquitously expressed with highest expression in mast cells and monocytes in human WAT, but in neutrophils and monocytes in mouse WAT (Figure 1C and D). CD44 was more abundantly expressed in subcutaneous WAT compared to omental WAT in humans (Figure 1E). In contrast, CD44 was more abundantly expressed in epididymal WAT relative to inguinal and periovarian WAT in mice (Figure 1F). Despite relatively low expression levels, CD44 was expressed in ASPCs (Figure 1C and D), which give rise to mature adipocytes and are essential for maintaining adipose tissue function. Furthermore, we compared the expression level of CD44 in epididymal WAT of mice fed with LFD (10% fat) or HFD (60% fat) using another single cell sequencing dataset (Sárvári et al., 2021). It was found that HFD feeding in mice increased the number of CD44 expressing cells, especially CD44 expressing immune cells, ASPCs and adipocytes (Fig 1G).

**Figure 1.**
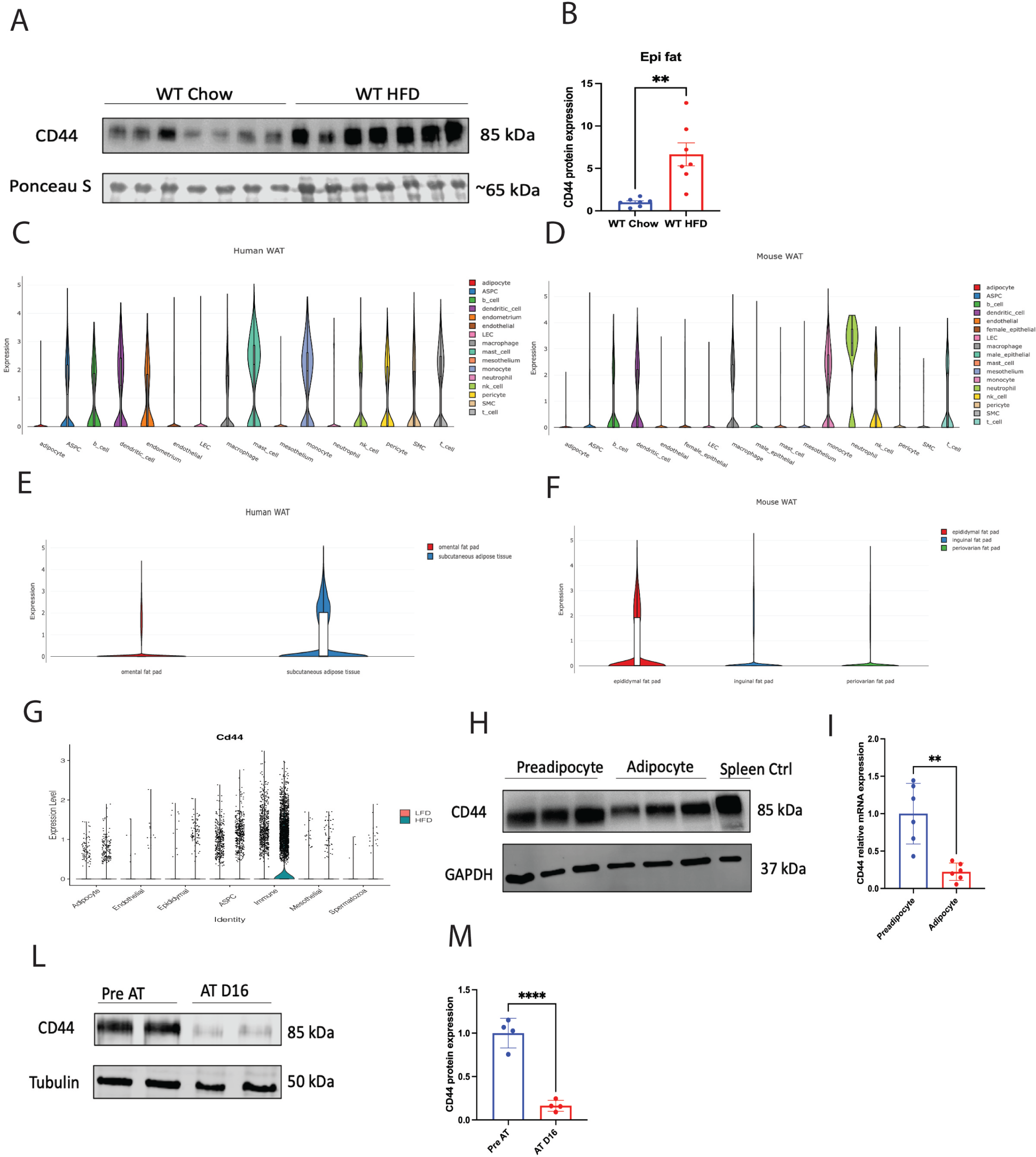
CD44 expression in adipose tissue, preadipocytes and adipocytes. (A-B) C57BL/6 mice were fed with either a chow diet (D/811004; DBM) or a 60% high fat diet (HFD) (#824054; SDS) for 16 weeks. CD44 protein expression was determined in the epididymal white adipose tissue (WAT) by Western. (C-D) CD44 clustering expression in the subcutaneous and visceral WAT of human and mice was analysed using published single cell RNA sequencing data (SCP1376) at Single Cell Portal (https://singlecell.broadinstitute.org/) (Emont et al., 2022). (E-F) CD44 expression in different depots of adipose tissue in human and mice was analysed using published single cell RNAseq data (SCP1376) at Single Cell Portal (https://singlecell.broadinstitute.org/) (Emont et al., 2022). (G) CD44 expression in different cell clusters in the epididymal WAT of male C57BL/6 mice fed 16 weeks of low-fat diet (LFD, 10% fat) or HFD diet (60% fat), was analysed using published single cell RNAseq data (SCP1179, https://singlecell.broadinstitute.org/) (Sárvári et al., 2021). (H-I) CD44 protein and mRNA expression in 3T3-L1 preadipocytes and 14-day differentiated adipocytes. (L-M) CD44 protein and mRNA expression in SGBS preadipocytes and 16-day differentiated adipocytes. Unpaired student t-test was performed for statistical comparison. \*\**p*<0.001, \*\*\*\**p*<0.0001.

In 3T3-L1 cells, both the mRNA and protein expression levels of CD44 were decreased after adipocyte differentiation (Figure 1H and I). Likewise, CD44 protein expression was down regulated in human Simpson-Golabi-Behmel syndrome (SGBS) adipocytes after differentiation (Figure 1L and M). These data suggest that CD44 was regulated in adipose tissue during obesity and expressed in preadipocytes, whose expression level was downregulated during adipocyte differentiation. Therefore, CD44 may have a role in regulating adipogenesis contributing to the pathogenesis of obesity.

### 3.2 Deletion of CD44 in pre-adipocytes promotes adipogenesis

To further study how CD44 regulates adipogenesis, we generated stable, CD44KO 3T3-L1 cells by Crispr Cas9-mediated gene editing. Three gRNAs targeting different exons of murine *Cd44* gene were designed (Supplemental Figure S1A) and constructed into Crispr Cas9 expressing vectors (PX459). The constructions were confirmed by enzymatic digestion and DNA sequencing (Supplemental Figure S1B and C). The 3T3-L1 pre-adipocytes were transfected by individual gRNA. The gRNA#3 transfected cells had most decreased CD44 expression when compared to other gRNAs and vector transfected cells (Supplemental Figure S1D), therefore were further sorted into single cell colonies. The stable CD44KO single cell lines were then screened for CD44 expression and two CD44KO cell lines (CD44KO1 and CD44KO2) with complete CD44 protein deletion were established (Figure 2A and Supplemental Figure S1E). Cells that underwent Crispr Cas9 editing but maintained normal CD44 protein expression were used as Crispr wildtype (WT) controls (Figure 2A). To further characterise the CD44KO1 and CD44KO2 cells, DNA sequences around the gRNA#3 targeted site were amplified from each single cell line (Supplemental Figure S1F) and bi-allelically sequenced. 3T3-L1 naïve WT cells and the Crispr WT cells had identical DNA sequence in both alleles of the *Cd44* gene (Figure 2B). However, CD44KO1 cells had two bases of AC deleted in both alleles (Figure 2C), and CD44KO2 cells had one base of A inserted in both alleles and a T to C mutation in the upstream of the gRNA#3 targeted site in one of the alleles (Figure 2D).

**Figure 2.**
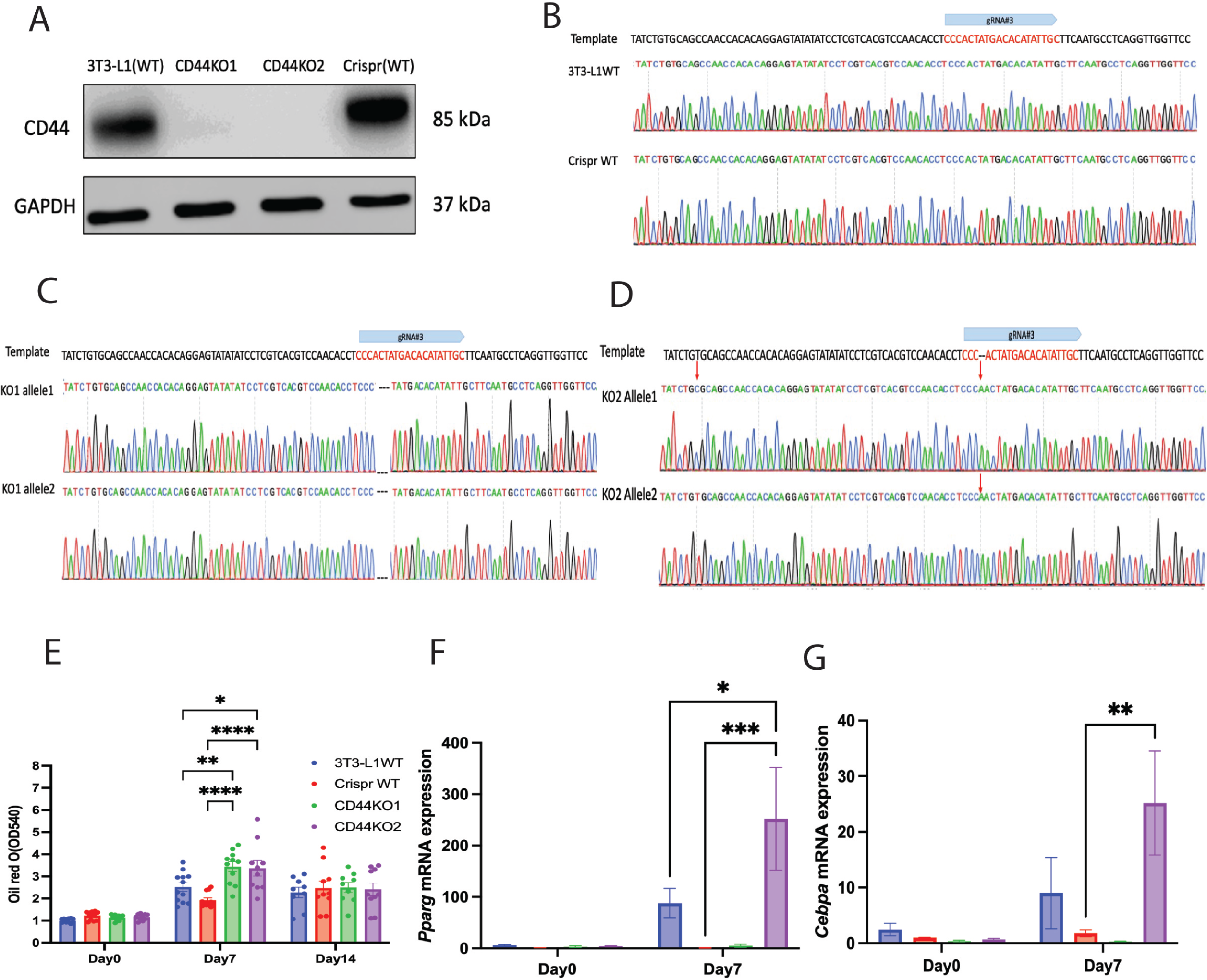
Deletion of CD44 promoted adipogenesis in 3T3-L1 cells. (A) Representative CD44 expression in 3T3-L1 naïve cells, Crispr WT and CD44KO cells. (B) Bi-allelic sequencing of 3T3-L1 naïve cells and Crispr WT cells. (C-D) Bi-allelic sequencing results of CD44KO1 and CD44KO2 cells. (E) Oil Red O staining during the differentiation of 3T3-L1 WT, Crispr WT and CD44KO cells. (F-G) *Pparg* and *Cebpa* mRNA expression on Day 0, and Day 7 after differentiation in 3T3-L1 WT, Crispr WT and CD44KO cells. Two-way ANOVA with Tukey’s multiple comparisons was performed to analyse the data. \**p*<0.05; \*\**p*<0.01; \*\*\**p*<0.005; \*\*\*\**p*<0.0001.

To determine how CD44 regulates adipogenesis, naïve 3T3-L1WT, CrisprWT, CD44KO1 and CD44KO2 cells were differentiated for 14 days. Adipogenesis was assessed by Oil Red O staining. The adipogenic induction significantly increased Oil Red O staining after 7 days of differentiation in all cells (Figure 2E). Interestingly, both CD44KO cell lines had significantly higher Oil Red O staining when compared to both 3T3-L1WT and CrisprWT cells (Figure 2E). These results suggest that deletion of CD44 promoted adipogenesis in 3T3-L1 cells. The expression of adipogenic transcriptional regulators *Pparg* and *Cebpa* were also measured before and 7 days after differentiation. While mRNA levels of *Pparg* and *Cebpa* were not different between naïve WT, CrisprWT and CD44KO1 cells, their expression levels were significantly increased in CD44KO2 cells on day 7 (Figure 2F and G). These results suggest that CD44KO1 and CD44KO2 cells promoted adipogenesis possibly via different mechanisms.

### 3.3 Lentivirus-mediated re-expression of CD44 attenuated adipogenesis in the CD44KO cells by suppressing *Pparg*

We next investigated CD44-specific effects on adipogenesis by re-expressing CD44 in the CD44KO cells. The expected molecular weight of the recombinant CD44 protein is 92KDa. CD44 re-expression (CD44RE) led to a 2-3-fold increase in CD44 protein expression at 92KDa in both CD44KO cell lines when compared to the empty vector transfected CD44KO cells (Figure 3A and B). CD44RE was also confirmed by the GFP expression detected at 92KDa. However, a GFP-tagged band at 65KDa was also observed in the CD44RE groups, which might correspond to degraded cytoplasmic fragments of CD44 protein. Consistent with the increased CD44 protein expression, CD44 mRNA levels were also increased in both CD44RE cell lines (Figure 3C). We next measured adipogenesis in these cells. Consistent with our hypothesis, CD44RE attenuated adipogenesis in both CD44KO cell lines with a 20% reduction in Oil Red O staining when compared to their respective CD44KO EV controls (Figure 3D and E). Because CD44KO2 cells promoted adipogenesis accompanied by upregulation of *Pparg* and *Cebpa* gene expression (Figure 2F and G), we further measured their expression in the CD44KO2 RE cells after 14 days of differentiation. Consistent with Oil Red O data, the *Pparg* mRNA expression was significantly decreased in the CD44KO2 RE cells when compared to CD44KO2 EV cells (Figure 3F). However, the mRNA expression of *Cebpa* was not changed between CD44KO2 RE and CD44KO2 EV cells (Figure 3G). These data suggest that CD44 could decrease *Pparg* expression, therefore inhibiting adipogenesis.

**Figure 3:**
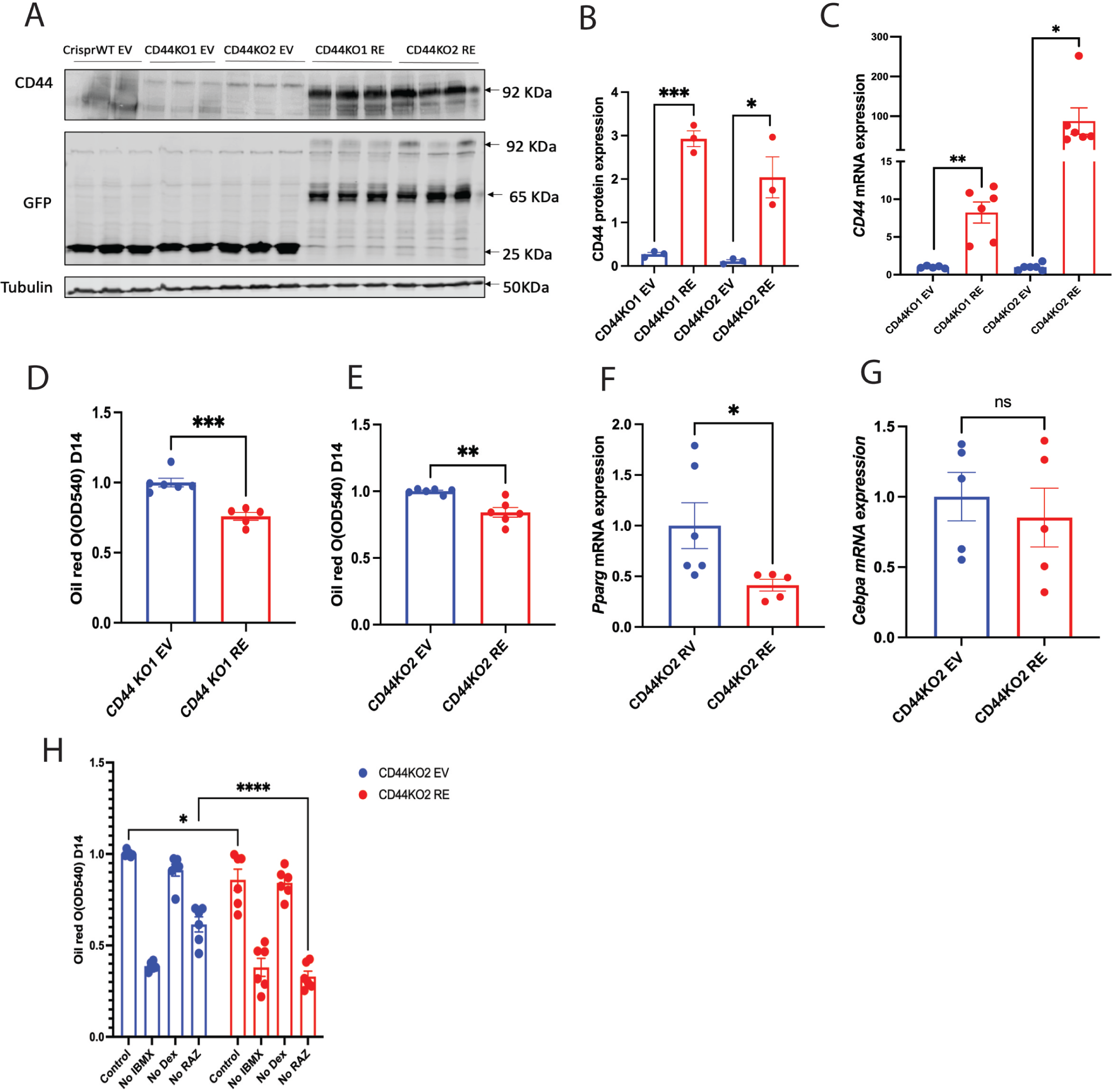
CD44RE attenuated adipogenesis via suppressing *Pparg* expression. (A) Representative Western blot of CD44 expression and GFP expression. (B) Quantification of CD44 protein expression. (C) mRNA expression of CD44 in CD44KO and CD44RE cells. (D-E) Oil Red O quantification on Day 14 of adipocytes differentiation in CD44KO and CD44RE cells. (F-G) *Pparg* and *Cebpa* mRNA expression in CD44KO2 EV and CD44KO2 RE cells after 14 days of differentiation. (H) CD44KO2 EV and CD44KO2 RE cells were differentiated for 14 days in adipogenic induction medium with/without specific agonists. IBMX, 3-isobutyl-1-methylxanthine; Dex, dexamethasone; RAZ, rosiglitazone. Unpaired student t-test or two-way ANOVA with Tukey’s multiple comparisons was performed for statistical analysis. \**p*<0.05; \*\**p*<0.01; \*\*\**p*<0.005; \*\*\*\**p*<0.0001.

To further confirm these results, we conditionally removed the agonists of various adipogenic stimuli in the differentiation medium of CD44KO2 EV and CD44KO2 RE cells. The cells were then differentiated for 7 days and adipogenesis was assessed. Consistently, CD44RE significantly decreased Oil Red O staining in the CD44KO2 cells when incubated with control differentiation cocktail (Figure 3H). Removal of IBMX, an activator of cAMP-associated pathway, significantly attenuated adipogenesis in both CD44KO2 EV and CD44KO2 RE cells, suggesting that activation of cAMP-associated signals was required for adipogenesis independent of CD44 expression. In contrast, removal of Dex, an activator of glucocorticoid receptor-associated pathway, had no effects on adipogenesis, indicating that CD44-mediated adipogenic regulation was unlikely through glucocorticoid receptor-associated pathways. Interestingly, the adipogenic cocktail without RAZ, a *Pparg* agonist attenuated adipogenesis in both CD44KO2 EV and CD44KO2 RE cells, when compared to the control cocktail, but to a much greater extend in the CD44KO2 RE cells when compared to CD44KO2 EV cells (Figure 3H). These results further demonstrate that CD44 could suppress *Pparg* expression, therefore inhibiting adipogenesis.

### 3.4 CD44RE in CD44KO cells impaired adipocyte function by decreasing insulin signalling and adiponectin secretion

To examine whether CD44RE attenuates adipocyte function, CD44KO EV and CD44KO RE cells were treated with 100nM insulin for 30mins after 14 days of differentiation. The insulin response was decreased by CD44RE in both CD44KO cell lines, evidenced by decreased AKT phosphorylation (S473) when compared to CD44KO EV cells (Figure 4A-D). In addition, adiponectin, a metabolically favourable hormone secreted by adipocytes, was decreased in differentiated CD44KO2 RE cells relative to CD44KO2 EV cells (Figure 4C and E), suggesting an impaired secretory/endocrine function of adipocytes.

**Figure 4:**
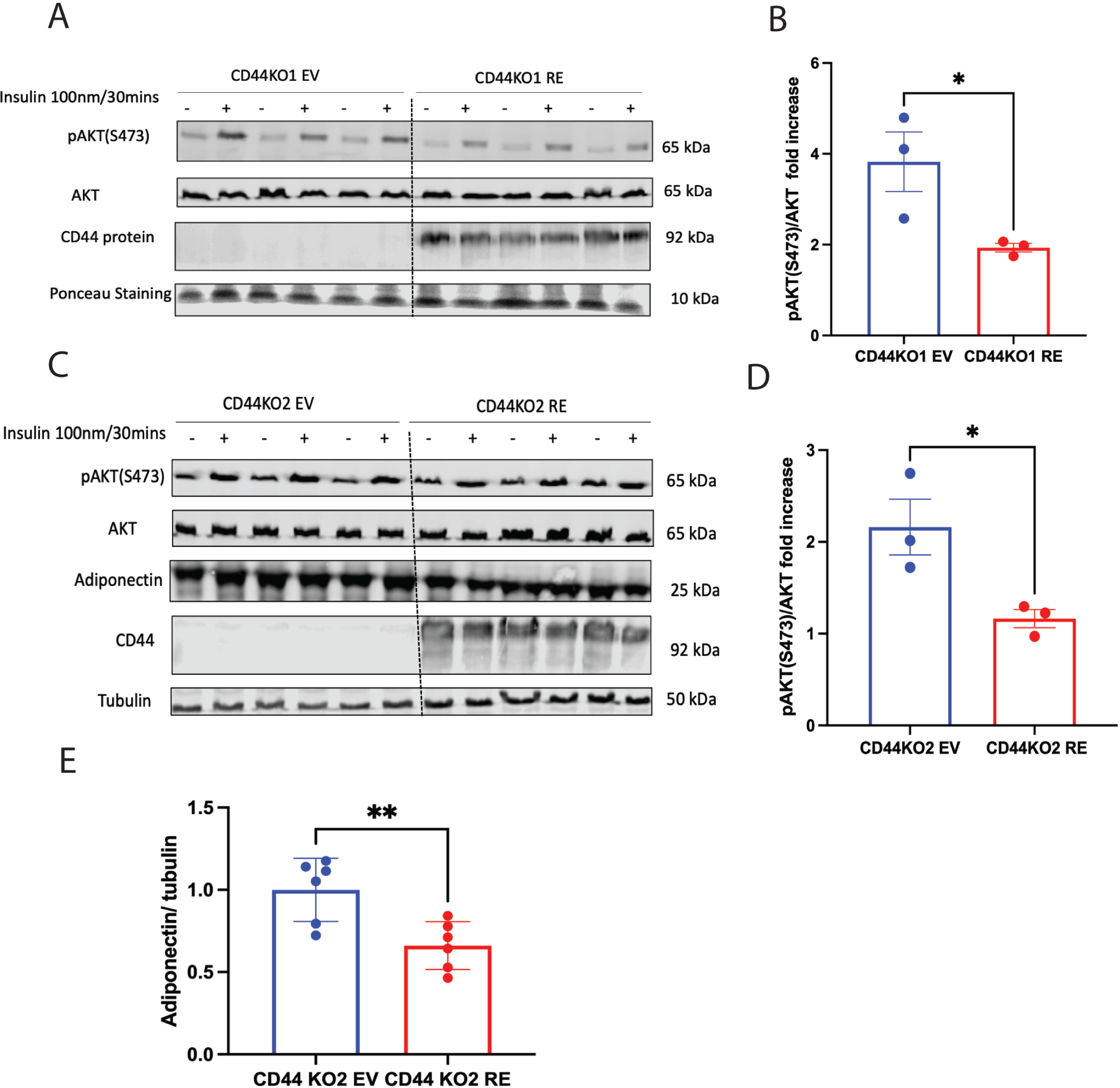
CD44RE decreased insulin responsiveness and adiponectin secretion. (A-B) CD44KO1 EV and CD44KO1 RE cells were differentiated for 14 days before AKT phosphorylation at S473 was measured with or without insulin stimulation. (C-D) CD44KO2 EV and CD44KO2 RE cells were differentiated for 14 days before AKT phosphorylation at S473 and adiponectin protein were measured with or without insulin stimulation. (E) Quantification of adiponectin expression. Unpaired student t-test was used for the statistical analysis. \**p*<0.05; \*\**p*<0.01.

### 3.5 Proteomics analysis of CrisprWT EV, CD44KO2 EV and CD44KO2 RE cells

Next, we applied proteomics to explore the potential mechanisms of how CD44 regulates adipogenesis and adipocyte function. To minimise the off-target effects of Crispr Cas9 gene editing, single cell sorting and lentiviral transfection, the CrisprWT EV cells were used as the control. CD44KO2 EV and CD44KO2 RE cells were studied due to their interesting phenotype on *Pparg*. Cells were validated by both CD44 protein and mRNA expression before the tandem mass tag (TMT)-based quantitative proteomics (Supplemental Figure 2A and B). Data were processed using Perseus (Supplemental Figure 2C). In total, 8541 proteins were identified from all three cell lines. A subset of 1087 DEPs were observed while comparing CD44KO2 EV to CrisprWT EV cells, with 548 proteins decreased and 539 proteins increased in the CD44KO2 EV cells. The proteomics results were visualized in volcano plots (Figure 5A and 5B), where the top 10 most-changed DEPs were marked. Surprisingly, between CD44KO2 RE and CD44KO2 EV cells only 30 DEPs were identified.

**Figure 5:**
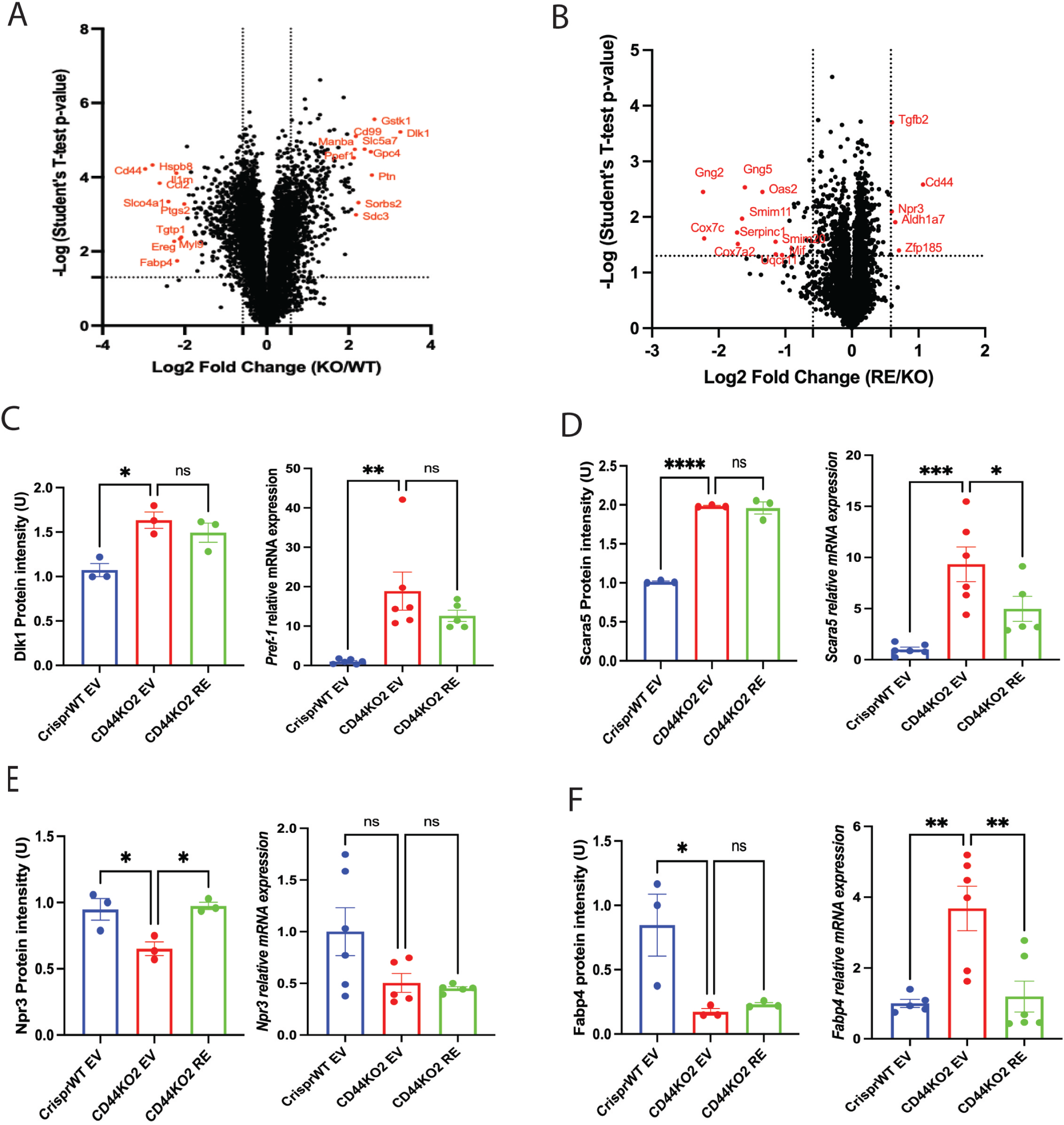
Proteomics analysis of CrisprWT EV, CD44KO2 EV and CD44KO2 RE cells. (A) Volcano plot showing differential protein expression in CD44KO2 EV vs CrisprWT EV cells. Data were plotted by Log2 fold change (KO/WT) to -Log(student t test *p* value). Top ten up and down regulated DEPs were marked in red. (B) Volcano plot showing differential protein expression in CD44KO2 RE vs CD44KO2 EV cells. Top ten down regulated DEPs, and five upregulated DEPs were marked in red. (C) Dlk1 protein expression and *Pref-1* mRNA expression. (D) Scara5 protein and mRNA expression. (E) Npr3 protein and mRNA expression. (F) Fabp4 protein and mRNA expression. Protein expression levels were determined by proteomics analysis and mRNA expression levels were measured by qRT-PCR. One-way ANOVA with Dunnett’s multiple comparisons was used for statistical analysis. ns: no statistical significance; \**p*<0.05; \*\**p*<0.01; \*\*\**p*<0.005; \*\*\*\**p*<0.0001.

We further explored the molecular regulation of CD44 on gene expression of some of the top 10 most-changed DEPs, which have been implicated in regulating adipogenesis. *Pref-1* (also known as *DlK1*) is a preadipocyte growth factor (Moon et al., 2002, Nueda et al., 2007), and was quantified as one of the mostly increased DEPs by the deletion of CD44 (Figure 5A). Consistent with the protein expression changes, its mRNA expression was increased by 20 folds in CD44KO2 EV when compared to CrisprWT EV cells (Figure 5C). However, there was no difference in *Pref-1* mRNA or protein expression between CD44KO2 EV and CD44KO2 RE cells (Figure 5C). *Scara5* is a scavenger receptor class A member protein, and previous study suggested that knockdown of *Scara5* inhibited adipogenesis in C3H10T1/2 pluripotent stem cells and A33 cells (Lee et al., 2017). Here, we found that deletion of CD44 significantly increased both the protein and mRNA expression of *Scara5,* while CD44RE attenuated its mRNA expression but not the protein expression (Figure 5D). *Npr3*, natriuretic peptide receptor 3, has been shown to arrest cell cycle progression (Li et al., 2021). We found that Npr3 was decreased after CD44 deletion and was one of the mostly increased DEPs by CD44RE (Figure 5B and E). However, mRNA level of *Npr3* was unchanged between CrisprWT EV, CD44KO2 EV and CD44KO2 RE cells (Figure 5E). Moreover, *Fabp4* is a fatty acid binding protein and identified as a marker of terminally differentiated adipocytes (Cao et al., 2013). Fabp4 protein was decreased after CD44 deletion (Figure 5F). However, its mRNA expression was significantly increased in CD44KO2 EV cells but was restored in CD44KO2 RE cells (Figure 5F).

The pathway enrichment was further analyzed using identified DEPs by metascape. Cell cycle-related pathways, such as mitotic cell cycle process, mitotic cell cycle phase transition, and cell cycle, were mostly enriched by CD44 deletion (Figure 6A). This implied that CD44 might regulate adipogenesis through regulating cell cycle progression. However, the pathway enrichment analysis between CD44KO2 RE and CD44KO2 EV cells was not possible due to small numbers of DEPs identified. Next, we analysed the protein expression of cell cycle related genes in the proteomics dataset. Deletion of CD44 caused a dramatic downregulation in proteins involved in all phases of cell cycle except for Pold4 (DNA polymerase delta 4) and Optn (optineurin) (Figure 6C). The downregulation in cell cycle-related proteins was consistent with the decrease in CD44 protein expression in CD44KO2 EV cells (Figure 6C). When CD44 was re-expressed in each cell line (CD44KO2 RE vs. CD44KO2 EV), this downregulation of most cell cycle-related proteins was rescued (Figure 6D). Altogether, our data suggest that CD44 may regulate adipogenesis via altering cell cycle progression.

**Figure 6.**
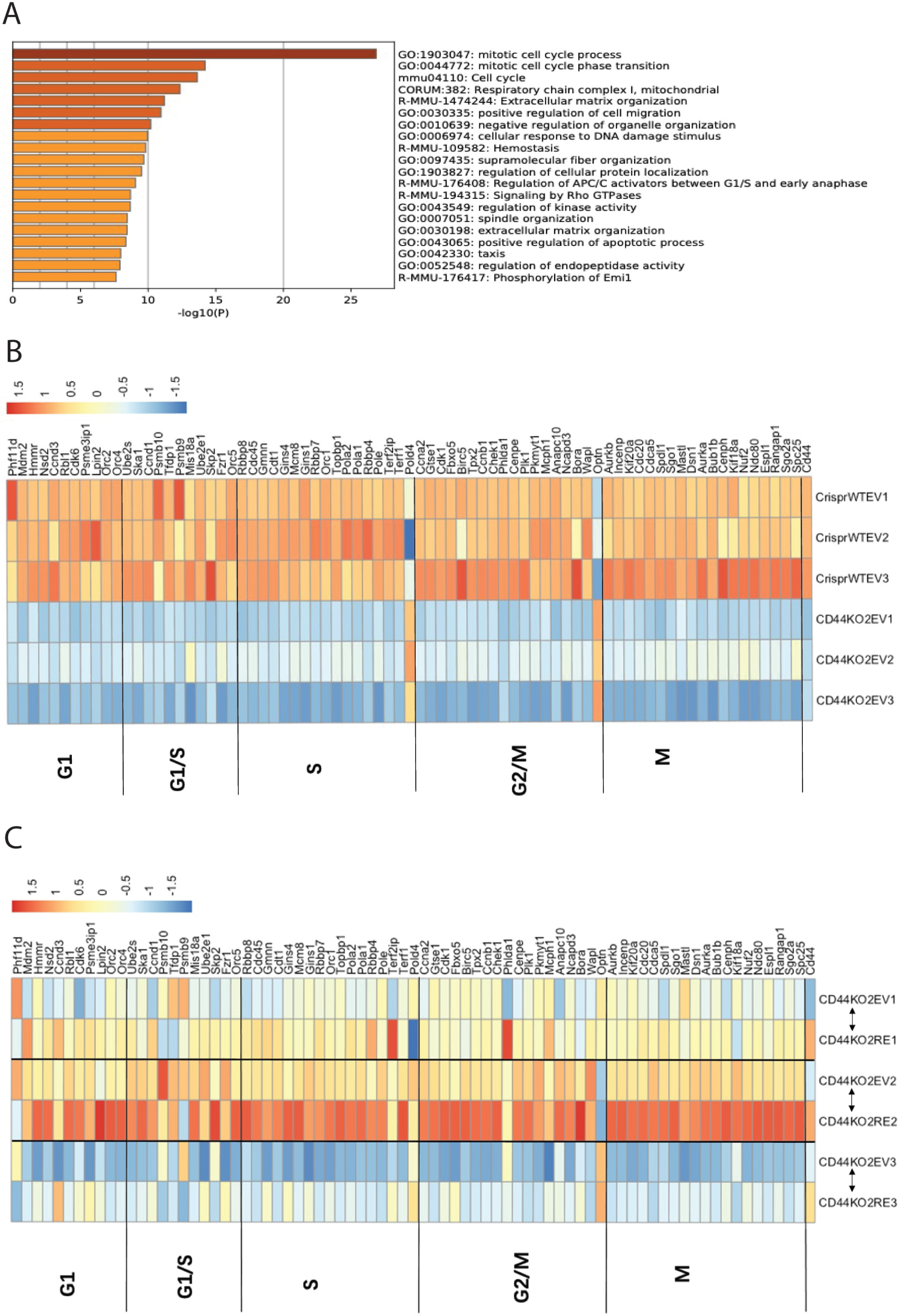
CD44 regulated adipogenesis possibly through cell cycle-related proteins. (A) Pathway enrichment analysis between CrisprWT EV vs CD44KO2 EV cells by metascape. (B-C) Heatmaps representing cell cycle-related protein expression in CrisprWT EV vs CD44KO2 EV, and CD44KO2 RE vs CD44KO2 EV with KEGG pathway enrichment cluster go:1903047, go:0044772 and mmu04110. Heatmaps were generated by pheatmap package in R (version 4.2.3).

## 4. Discussion

CD44, a well-known stem cell biomarker is recently implicated in chronic metabolic diseases (Weng et al., 2022). CD44 and its ligands HA and OPN were increased in the adipose tissue of obese mice (Kang et al., 2013, Kiefer et al., 2008, Nomiyama et al., 2007, Zhu et al., 2021, Romo et al., 2022). Here, we found for the first time that CD44 expression in preadipocytes hinders adipocyte differentiation and adipogenesis, therefore negatively impacting adipocyte function including insulin responsiveness and its secretory capacity. Mechanistically, CD44 does so possibly through suppressing adipogenic transcription factor *Pparg* expression and cell cycle-related pathways.

CD44 has previously shown to be primarily expressed in inflammatory cells such as macrophages in obese adipose tissue and CD44 expression positively correlated with inflammatory markers CD68 and IL6 gene expression in the subcutaneous WAT of obese human (Kodama et al., 2012, Liu et al., 2015). In consistent with these findings, we observed that CD44 was highly expressed in mast cells and monocytes in human WAT, and monocytes and neutrophils in mouse WAT. But a considerable amount of CD44 was also expressed in ASPCs in human WAT. Moreover, in obese mice, in addition to immune cells CD44 expressing cells mainly enriched in ASPC cells and adipocytes. These results support the notion that CD44 may be important in regulating adipose function through its direct action in adipocytes or adipocyte precursor cells. It is worth noting that CD44 was differentially expressed in human and mouse WAT, with the highest expression in subcutaneous WAT in humans but in epididymal WAT in mice. These data suggest a potentially differential role of CD44 in human and mice since subcutaneous adipose tissue in human is considered metabolically beneficial while epididymal adipose tissue in mice is detrimental to metabolic regulation during obesity (Wajchenberg et al., 2002, Boyko et al., 2000, Misra et al., 1997, Snijder et al., 2004). The species differences of CD44 expression in adipose tissue remains unknown and will merit future investigations.

Adipose *Cd44* gene was associated with T2D by an expression based-genome wide association study (Kodama et al., 2012), however, it is unclear which mutations in the coding region of *Cd44* or which CD44 variant may be involved. Using CRISPR Cas9-mediated gene editing technique, we established two stable CD44KO cell lines, which adopted distinct genetic decorations to ablate CD44. Interestingly, despite the distinct genetic manipulations, both KO cells promoted adipogenesis in 3T3-L1 cells. However, this promotion was only observed 7 days after differentiation initiation but was lost by Day 14. This suggests that CD44 may be more crucial in regulating mitotic proliferation and cell cycle arrest than terminal differentiation and maturation of adipocytes (Chang and Kim, 2019). Moreover, the two KO cell lines enhanced early differentiation through different mechanisms, where CD44 KO2 but not CD44 KO1 cells had increased *Pparg* and *Cebpa* expression relative to 3T3-L1WT and CrisprWT cells on Day 7. The discrepant regulation of *Pparg* and *Cebpa* expression in two CD44KO cell lines may be due to differences in genetic mutations (bp deletion in CD44KO1 vs. insertion in CD44KO2). Indeed, genetic deletion was suggested to cause more harm than insertion mutation because a deleted DNA fragment is difficult to be replaced at the exact position (Pineda et al., 2019). Regardless, the gene expression of *Pparg* and *Cebpa* were not induced after 7 days of differentiation in the CrisprWT cells unlike in the 3T3-L1 naïve cells, which may suggest potential off-target effects of CRISPR Cas9 gene editing. While the gRNAs guide the Cas9 protein to break DNA sequences of interest in the genome, it can also promote Cas9 protein to mismatch targeted DNA sequences (Zhang et al., 2015), resulting in off-target effects. However, if any, these off-target effects will be equivalent between the CrisprWT control cells and the two KO cell lines.

Here, we observe an inhibitory role of CD44 in adipogenesis, at least in 3T3-L1 cells, as deletion of CD44 promoted lipid accumulation during adipocyte differentiation and re-expression of CD44 in CD44 KO cells abolished this effect, accentuating CD44 specific effects. Kang et al., reported that HFD-fed CD44 global KO mice had increased epididymal WAT mass with larger adipocytes and increased lipogenic gene expression (e.g. *Cidec, Fasn, Fabp1, Mogat2*) (Kang et al., 2013), indicative of enhanced adipogenesis and lipogenesis, which support our findings. Moreover, we provided additional mechanistic evidence how CD44 may regulate adipogenesis via modulating *Pparg* expression. Activation of *Pparg* by rosiglitazone has been previously shown to promote marrow mesenchymal stem cell U-33/γ2 differentiation with decreased CD44 (Shockley et al., 2007), suggesting a possible negative regulation between *Pparg* and CD44. Furthermore, our conditional culture medium studies confirmed that lack of *Pparg* activation significantly attenuated adipogenesis in both CD44KO and CD44RE cells, but to a much greater extent in the CD44RE cells. Taken together, these results suggest that CD44 suppressed *Pparg* expression, therefore, inhibiting adipogenesis.

We further explored potential mechanisms by which CD44 inhibits adipogenesis through unbiased proteomics analysis. The metascope pathway enrichment analysis suggests that cell cycle progression-related pathways, e.g. mitotic cell cycle process and mitotic cell phase transition were most changed by deletion of CD44 in preadipocytes. Preadipocytes undergo unlimited proliferation until they become growth-arrested through contact inhibition (Lefterova and Lazar, 2009). Under the stimulation of hormones such as insulin, the growth-arrested preadipocytes undergo cell cycle re-entry for mitotic clonal expansion and subsequent adipogenic induction (Lefterova and Lazar, 2009). Interestingly, our proteomics data suggest that most proteins involved in all phases of the cell cycle were decreased by deletion of CD44 and was restored by re-expression of CD44. These results suggest that deletion of CD44 in preadipocytes may decrease cell division but increase growth arrest, therefore inducing cells towards mitotic clonal expansion and lipid accumulation. However, this hypothesis remains to be tested. It is important to recognise that overall changes in protein signatures caused by CD44 re-expression were moderate. This could be due to insufficient re-expression of CD44 (Supplemental Figure S2A-B) or possibly fragmented CD44 as identified by a 65KDa GFP-tagged protein band in Western blotting (Figure 3A). Other protein expression strategies including direct transfer of plasmid DNA of CD44 are needed to further uncover CD44 specific proteomic effects.

In addition, we also observed some interesting regulation of CD44 on the mRNA and protein expression of various factors that have been shown to regulate adipogenesis, including Pref-1, Scara5, Npr3, and Fabp4 (Moon et al., 2002, Nueda et al., 2007, Lee et al., 2017, Li et al., 2021, Cao et al., 2013). Particularly, gene expression of *Scara5*, a novel mediator of adipocyte commitment, was increased by deletion of CD44, which was partially rescued by re-expression of CD44. Protein expression of Npr3, an atrial natriuretic peptide receptor which was recently shown to inhibit insulin-stimulated glucose uptake and *de novo* lipogenesis in adipocytes (Roos et al., 2021), was decreased by deletion of CD44 and restored by re-expression of CD44. These results suggest that CD44 may regulate adipogenesis and adipocyte function through other novel mediators, which merits further investigations.

Blocking CD44 by genetic ablation or antibody antagonism attenuates adipose tissue inflammation, fibrosis, and insulin resistance with increased adiposity in obese mice (Kodama et al., 2015, Kang et al., 2013). Genetic deletion of CD44 also ameliorates HFD-induced skeletal muscle insulin resistance in obese mice (Hasib et al., 2019). CD44 abrogates insulin responsiveness upon activation by its ligands HA and OPN (Weng et al., 2022). Here, we found that re-expression of CD44 in CD44KO adipocytes decreased the insulin response by downregulating AKT phosphorylation (S473) and decreased adiponectin secretion. Adipose tissue stores excess energy in the form of triglycerides via *de novo* lipogenesis, adipogenesis and adipocyte hypertrophy during obesity. Impaired adipogenesis pressurises adipocytes into hypertrophy and subsequent adipocyte death, contributing to insulin resistance *in vivo* (Gustafson et al., 2015, Vishvanath and Gupta, 2019). Thus, the regulation of CD44 on adipogenesis in preadipocytes may be crucial for determining insulin responsiveness and endocrine functions of mature adipocytes. Regardless, it is important to recognise that CD44 is primarily expressed in immune cells which were not tested in the current study but remains an important contributor to adipose function.

In conclusion, our study utilised CRISPR Cas9-mediated *Cd44* deletion and lentivirus-mediated *Cd44* re-expression techniques in 3T3-L1 cells and discovered that CD44 in preadipocytes is a critical regulator of adipocyte differentiation and the endocrine function of mature adipocytes. We are the first to study the direct role of (pre)adipocyte CD44 in adipose function, providing insight into a potential extracellular matrix-receptor regulatory component, the HA/OPN-CD44 signalling in adipocytes. It is currently unknown whether the favourably metabolic effects of CD44 deletion in obesity is through cell-autonomous actions or non-adipocytes e.g. macrophages and adipocytes interactions. Future studies using CD44 conditional KO mice to investigate cell type specific role of CD44 in metabolism is essential. Nevertheless, our studies together with previous evidence suggest that CD44 may be a therapeutic target to treat obesity associated metabolic diseases such as T2D.

## Author Contributions

XW and LK contributed to the experimental design, researched data, contributed to discussion and data interpretation, and wrote the manuscript. HJ, HZ, DL, and JW researched data, and reviewed and edited the manuscript.

## Acknowledgement

LC-MS analysis was done at the FingerPrints Proteomics Facility at the University of Dundee.

## References

Boyko, E. J., Fujimoto, W. Y., Leonetti, D. L. & Newell-Morris, L. 2000. Visceral adiposity and risk of type 2 diabetes: a prospective study among Japanese Americans. Diabetes Care, 23, 465–71.

Cao, H., Sekiya, M., Ertunc, M. E., Burak, M. F., Mayers, J. R., White, A., Inouye, K., Rickey, L. M., Ercal, B. C., Furuhashi, M., Tuncman, G. & Hotamisligil, G. S. 2013. Adipocyte lipid chaperone AP2 is a secreted adipokine regulating hepatic glucose production. Cell Metab, 17, 768–78.

Chang, E. & Kim, C. Y. 2019. Natural Products and Obesity: A Focus on the Regulation of Mitotic Clonal Expansion during Adipogenesis. Molecules, 24.

Collaboration, N. C. D. R. F. 2017. Worldwide trends in body-mass index, underweight, overweight, and obesity from 1975 to 2016: a pooled analysis of 2416 population-based measurement studies in 128.9 million children, adolescents, and adults. Lancet, 390, 2627–2642.

Emont, M. P., Jacobs, C., Essene, A. L., Pant, D., Tenen, D., Colleluori, G., DI Vincenzo, A., Jørgensen, A. M., Dashti, H., Stefek, A., Mcgonagle, E., Strobel, S., Laber, S., Agrawal, S., Westcott, G. P., Kar, A., Veregge, M. L., Gulko, A., Srinivasan, H., Kramer, Z., DE Filippis, E., Merkel, E., Ducie, J., Boyd, C. G., Gourash, W., Courcoulas, A., Lin, S. J., Lee, B. T., Morris, D., Tobias, A., Khera, A. V., Claussnitzer, M., Pers, T. H., Giordano, A., Ashenberg, O., Regev, A., Tsai, L. T. & Rosen, E. D. 2022. A single-cell atlas of human and mouse white adipose tissue. Nature, 603, 926–933.

Gustafson, B., Hedjazifar, S., Gogg, S., Hammarstedt, A. & Smith, U. 2015. Insulin resistance and impaired adipogenesis. Trends Endocrinol Metab, 26, 193–200.

Hasib, A., Hennayake, C. K., Bracy, D. P., BUGLER-Lamb, A. R., Lantier, L., Khan, F., Ashford, M. L. J., Mccrimmon, R. J., Wasserman, D. H. & Kang, L. 2019. CD44 contributes to hyaluronan-mediated insulin resistance in skeletal muscle of high-fat-fed C57BL/6 mice. Am J Physiol Endocrinol Metab, 317, E973–E983.

HASSN Mesrati, M., Syafruddin, S. E., Mohtar, M. A. & Syahir, A. 2021. CD44: A Multifunctional Mediator of Cancer Progression. Biomolecules, 11.

Hughes, C. S., Moggridge, S., Müller, T., Sorensen, P. H., Morin, G. B. & Krijgsveld, J. 2019. Single-pot, solid-phase-enhanced sample preparation for proteomics experiments. Nat Protoc, 14, 68–85.

Kang, H. S., Liao, G., Degraff, L. M., Gerrish, K., Bortner, C. D., Garantziotis, S. & Jetten, A. M. 2013. CD44 plays a critical role in regulating diet-induced adipose inflammation, hepatic steatosis, and insulin resistance. PLoS One, 8, e58417.

Kiefer, F. W., Zeyda, M., Todoric, J., Huber, J., Geyeregger, R., Weichhart, T., Aszmann, O., Ludvik, B., Silberhumer, G. R., Prager, G. & Stulnig, T. M. 2008. Osteopontin expression in human and murine obesity: extensive local up-regulation in adipose tissue but minimal systemic alterations. Endocrinology, 149, 1350–7.

Klein, S., Gastaldelli, A., YKI-Järvinen, H. & Scherer, P. E. 2022. Why does obesity cause diabetes? Cell Metab, 34, 11–20.

Kodama, K., Horikoshi, M., Toda, K., Yamada, S., Hara, K., Irie, J., Sirota, M., Morgan, A. A., Chen, R., Ohtsu, H., Maeda, S., Kadowaki, T. & Butte, A. J. 2012. Expression-based genome-wide association study links the receptor CD44 in adipose tissue with type 2 diabetes. Proc Natl Acad Sci U S A, 109, 7049–54.

Kodama, K., Toda, K., Morinaga, S., Yamada, S. & Butte, A. J. 2015. Anti-CD44 antibody treatment lowers hyperglycemia and improves insulin resistance, adipose inflammation, and hepatic steatosis in diet-induced obese mice. Diabetes, 64, 867–75.

Lee, H., Lee, Y. J., Choi, H., Seok, J. W., Yoon, B. K., Kim, D., Han, J. Y., Lee, Y., Kim, H. J. & Kim, J. W. 2017. SCARA5 plays a critical role in the commitment of mesenchymal stem cells to adipogenesis. Sci Rep, 7, 14833.

Lefterova, M. I. & Lazar, M. A. 2009. New developments in adipogenesis. Trends Endocrinol Metab, 20, 107–14.

Li, S., Guo, R., Peng, Z., Quan, B., Hu, Y., Wang, Y. & Wang, Y. 2021. NPR3, transcriptionally regulated by POU2F1, inhibits osteosarcoma cell growth through blocking the PI3K/AKT pathway. Cell Signal, 86, 110074.

Liu, L. F., Kodama, K., Wei, K., Tolentino, L. L., Choi, O., Engleman, E. G., Butte, A. J. & Mclaughlin, T. 2015. The receptor CD44 is associated with systemic insulin resistance and proinflammatory macrophages in human adipose tissue. Diabetologia, 58, 1579–86.

Misra, A., Garg, A., Abate, N., Peshock, R. M., STRAY-Gundersen, J. & Grundy, S. M. 1997. Relationship of anterior and posterior subcutaneous abdominal fat to insulin sensitivity in nondiabetic men. Obes Res, 5, 93–9.

Moon, Y. S., Smas, C. M., Lee, K., Villena, J. A., Kim, K. H., Yun, E. J. & Sul, H. S. 2002. Mice lacking paternally expressed Pref-1/Dlk1 display growth retardation and accelerated adiposity. Mol Cell Biol, 22, 5585–92.

Nomiyama, T., PEREZ-Tilve, D., Ogawa, D., Gizard, F., Zhao, Y., Heywood, E. B., Jones, K. L., Kawamori, R., Cassis, L. A., Tschöp, M. H. & Bruemmer, D. 2007. Osteopontin mediates obesity-induced adipose tissue macrophage infiltration and insulin resistance in mice. J Clin Invest, 117, 2877–88.

Nueda, M. L., Baladron, V., SANCHEZ-Solana, B., Ballesteros, M. A. & Laborda, J. 2007. The EGF-like protein dlk1 inhibits notch signaling and potentiates adipogenesis of mesenchymal cells. J Mol Biol, 367, 1281–93.

Pineda, M., Lear, A., Collins, J. P. & Kiani, S. 2019. Safe CRISPR: Challenges and Possible Solutions. Trends Biotechnol, 37, 389–401.

Ponta, H., Sherman, L. & Herrlich, P. A. 2003. CD44: from adhesion molecules to signalling regulators. Nat Rev Mol Cell Biol, 4, 33–45.

Romo, M., López-Vicario, C., Pérez-Romero, N., Casulleras, M., Martínez-Puchol, A. I., Sánchez, B., Flores-Costa, R., Alcaraz-Quiles, J., Duran-Güell, M., Ibarzábal, A., Espert, J. J., Clària, J. & Titos, E. 2022. Small fragments of hyaluronan are increased in individuals with obesity and contribute to low-grade inflammation through TLR-mediated activation of innate immune cells. Int J Obes (Lond), 46, 1960–1969.

Roos, J., Dahlhaus, M., Funcke, J. B., Kustermann, M., Strauss, G., Halbgebauer, D., Boldrin, E., Holzmann, K., Moller, P., Trojanowski, B. M., Baumann, B., Debatin, K. M., Wabitsch, M. & Fischer-Posovszky, P. 2021. miR-146a regulates insulin sensitivity via NPR3. Cell Mol Life Sci, 78, 2987–3003.

Sárvári, A. K., VAN Hauwaert, E. L., Markussen, L. K., Gammelmark, E., Marcher, A. B., Ebbesen, M. F., Nielsen, R., Brewer, J. R., Madsen, J. G. S. & Mandrup, S. 2021. Plasticity of Epididymal Adipose Tissue in Response to Diet-Induced Obesity at Single-Nucleus Resolution. Cell Metab, 33, 437–453.e5.

Shockley, K. R., Rosen, C. J., Churchill, G. A. & Lecka-Czernik, B. 2007. PPARgamma2 Regulates a Molecular Signature of Marrow Mesenchymal Stem Cells. PPAR Res, 2007, 81219.

Snijder, M. B., Dekker, J. M., Visser, M., Bouter, L. M., Stehouwer, C. D., Yudkin, J. S., Heine, R. J., Nijpels, G., Seidell, J. C. & Hoorn, S. 2004. Trunk fat and leg fat have independent and opposite associations with fasting and postload glucose levels: the Hoorn study. Diabetes Care, 27, 372–7.

Stefan, N. 2020. Causes, consequences, and treatment of metabolically unhealthy fat distribution. Lancet Diabetes Endocrinol, 8, 616–627.

Vishvanath, L. & Gupta, R. K. 2019. Contribution of adipogenesis to healthy adipose tissue expansion in obesity. J Clin Invest, 129, 4022–4031.

Wajchenberg, B. L., Giannella-Neto, D., Da Silva, M. E. & Santos, R. F. 2002. Depot-specific hormonal characteristics of subcutaneous and visceral adipose tissue and their relation to the metabolic syndrome. Horm Metab Res, 34, 616–21.

Weng, X., Maxwell-Warburton, S., Hasib, A., Ma, L. & Kang, L. 2022. The membrane receptor CD44: novel insights into metabolism. Trends Endocrinol Metab, 33, 318–332.

Zhang, X. H., Tee, L. Y., Wang, X. G., Huang, Q. S. & Yang, S. H. 2015. Off-target Effects in CRISPR/Cas9-mediated Genome Engineering. Mol Ther Nucleic Acids, 4, e264.

Zhu, Y., Li, N., Huang, M., Bartels, M., Dogné, S., Zhao, S., Chen, X., Crewe, C., Straub, L., Vishvanath, L., Zhang, Z., Shao, M., Yang, Y., Gliniak, C. M., Gordillo, R., Smith, G. I., Holland, W. L., Gupta, R. K., Dong, B., Caron, N., Xu, Y., Akgul, Y., Klein, S. & Scherer, P. E. 2021. Adipose tissue hyaluronan production improves systemic glucose homeostasis and primes adipocytes for CL 316,243-stimulated lipolysis. Nat Commun, 12, 4829.

Zöller, M. 2011. CD44: can a cancer-initiating cell profit from an abundantly expressed molecule? Nat Rev Cancer, 11, 254–67.

