## Supplemental materials for "Deletion of CD44 promotes adipogenesis by regulating PPARɣ and cell cycle-related pathways"

**LC-MS analysis**

LC buffers were the following:  buffer A (0.1% formic acid in Milli-Q water (v/v)) and buffer B (80% acetonitrile and 0.08% formic acid in Milli-Q water (v/v). Aliquots of 2μL of each sample were loaded at 10μL/min onto a trap column (100μm × 2cm, PepMap nanoViper C18 column, 5μm, 100Å) equilibrated in 5% buffer B. The trap column was washed for 5min at the same flow rate and then the trap column was switched in-line with a Thermo Scientific, resolving C18 column (75μm × 50cm, PepMap RSLC C18 column, 2μm, 100Å). The peptides were eluted from the column at a constant flow rate of 300nL/min with a linear gradient from 5% buffer B (for Fractions 1-10, 7% for Fractions 11-20) to 35% buffer B in 130min, and then to 98% buffer B by 132min. The column was then washed with 98% buffer B for 20min and re-equilibrated in 5% or 7% buffer B for 17min. The MS spectra was acquired using data dependent mode (DDA). A scan cycle comprised MS1 scan (m/z range from 335-1800, with a maximum ion injection time of 50ms, a resolution of 120,000 and automatic gain control (AGC) value of 3x10^6^) followed by 15 sequential dependant MS2 scans (with an isolation window set to 0.7m/z, resolution at 60,000, maximum ion injection time at 200ms and AGC 1x10^5^). To ensure mass accuracy, the mass spectrometer was calibrated on the first day that the runs were performed.

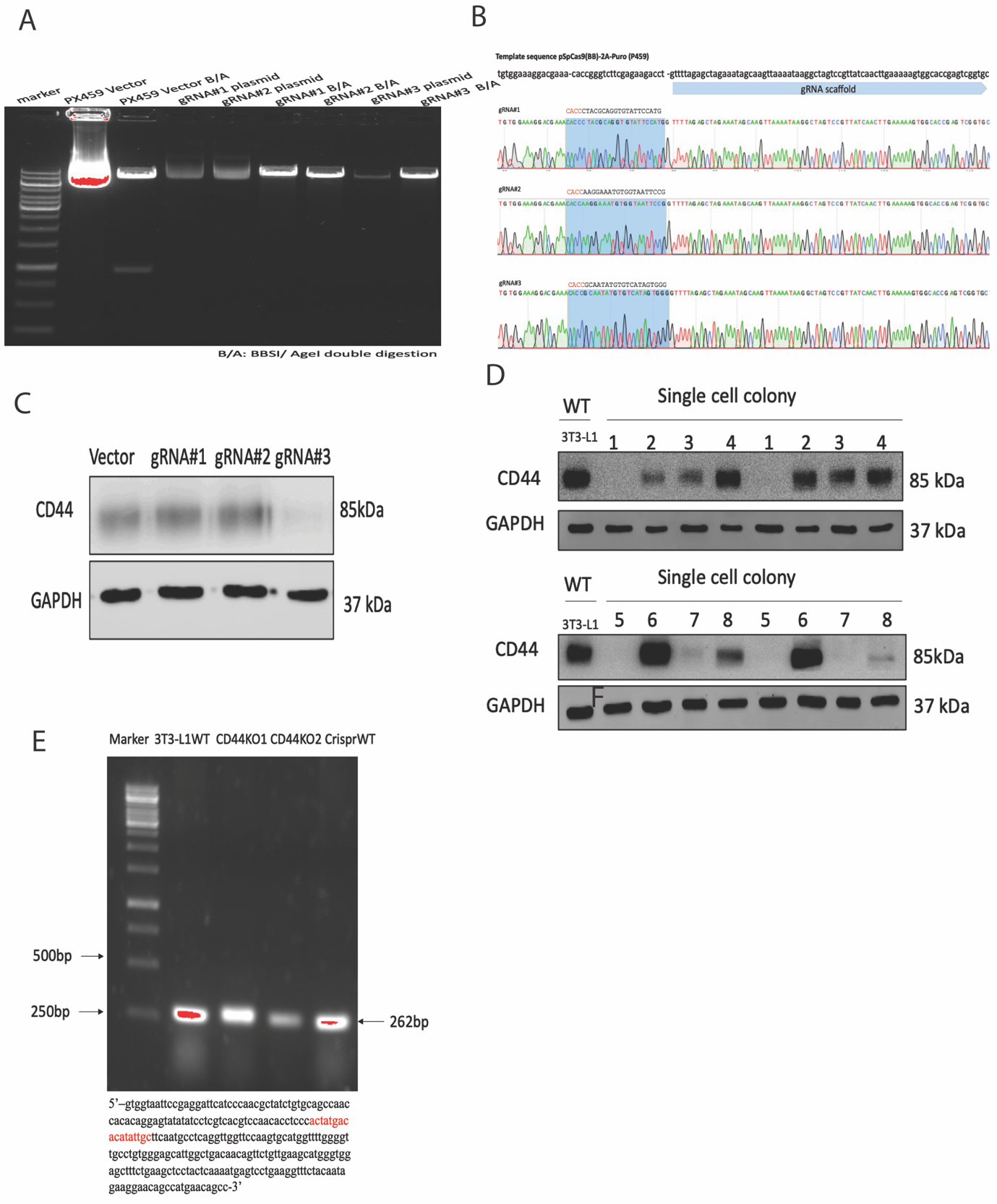

**Figure supplement 1: Generation of CD44KO 3T3-L1 cells using Crispr Cas9 gene editing.** (A) gRNA design using the Broad institute web portal (<http://www.broadinstitute.org/rnai/public/analysis-tools/sgrna-design>). (B) BbsI and AgeI double enzyme digestion of gRNA containing PX459 plasmid. (C) Sequencing confirmation of the insertion of gRNA in PX459 plasmid. (D) CD44 protein expression after gRNA transfection. (E) Stable CD44KO single cell sequencing after sorting. (F) PCR amplification of the DNA fragment of *Cd44* gene containing gRNA#3 edited sequence.

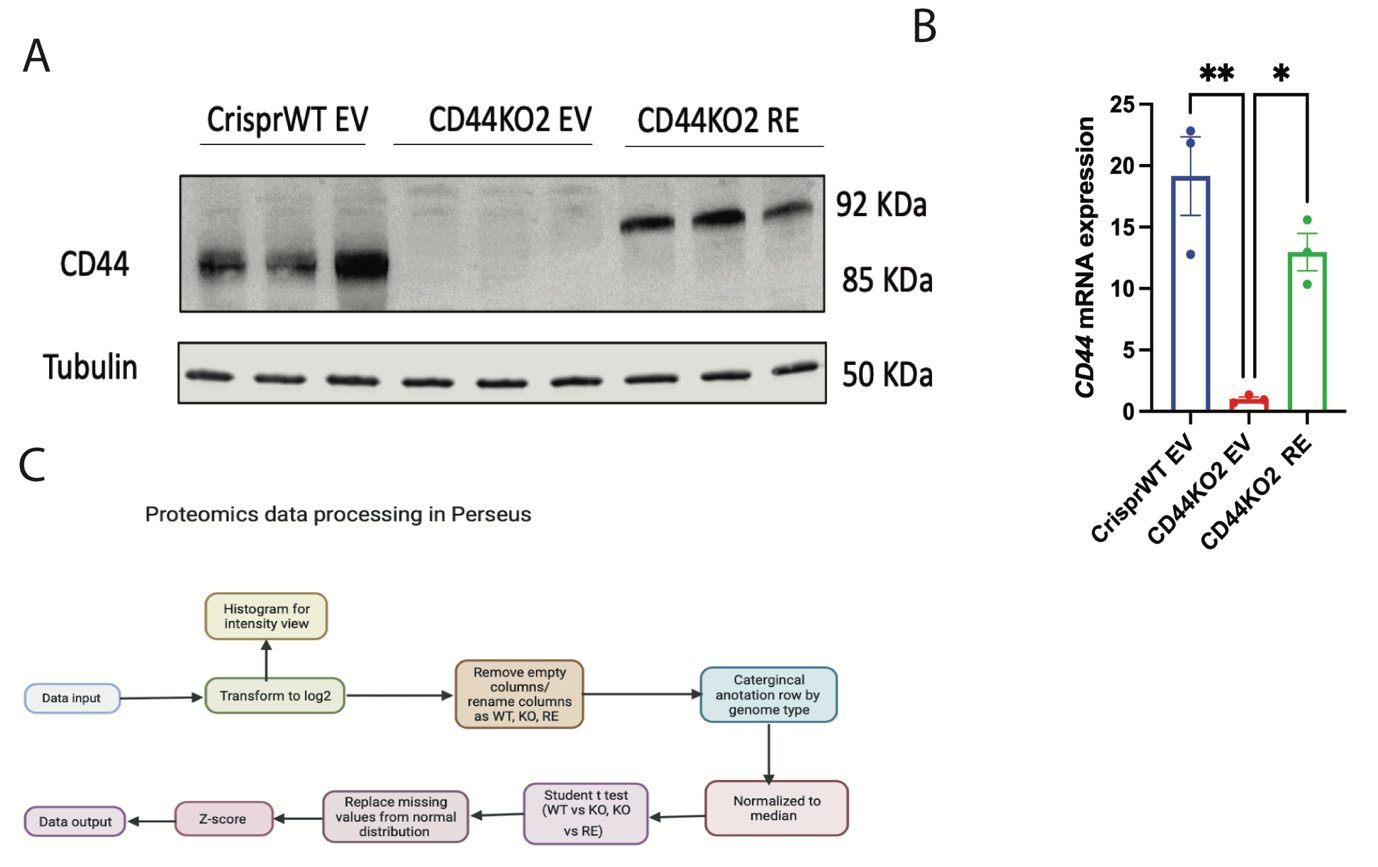

**Figure supplement 2: Proteomics analysis of CrisprWT EV, CD44KO2 EV and CD44KO2 RE cells.** (A) and (B) CD44 protein and mRNA expression validation before proteomics analysis. (C) Workflow of proteomics data processing in Perseus software.
